# Adjuvants MPLA and SMNP induce antiviral immunity and indirectly revert HIV-1 latency

**DOI:** 10.64898/2026.04.24.720763

**Authors:** Jade Jansen, Killian E. Vlaming, Leanne C. Helgers, Eva Schumacher, Tanja M. Kaptein, Rogier W. Sanders, Godelieve J. de Bree, Teunis B.H. Geijtenbeek, Neeltje A. Kootstra

## Abstract

Monophosphoryl lipid A (MPLA) and the saponin-MPLA nanoparticle adjuvant (SMNP) are known immunostimulants used as adjuvants in vaccines to boost immunity. These adjuvants are under investigation for use in prophylactic and therapeutic HIV-1 vaccines. However, their effects in ART-treated people with HIV-1 (PWH) remain unclear, as chronic immune activation may reduce responses and potentially reactivate latent virus reservoirs.

Here we observed that both adjuvants, MPLA and SMNP, triggered TLR4-dependent cytokine production in monocyte-derived dendritic cells (DCs), but SMNP elicited a stronger response than MPLA based on costimulatory receptor expression and cytokine production. Cytokines produced by adjuvant stimulated dendritic cells were able to induce HIV-1 transcription in a J-Lat cell model.

Notably, SMNP, but not MPLA, also induced a cytokine response in PBMC from PWH, albeit lower as compared to in healthy donor PBMC. Importantly, SMNP substantially reduced the size of the inducible HIV-1 reservoir in PWH *ex vivo* without bystander cytotoxicity. The effect on the viral reservoir was likely caused by the cytokine production induced by SMNP, as supernatants from SMNP stimulated DCs showed a similar viral reservoir reduction.

Our findings demonstrate that TLR4-targeted adjuvants, especially SMNP, can effectively induce immune activation, supporting their potential use in therapeutic vaccination. However, as reservoir reduction was observed *ex vivo*, further evaluation is warranted to ensure safe use of such adjuvants in PWH.

## Introduction

Adjuvants are important in vaccine strategies to induce innate immunity and thereby elicit strong adaptive immunity. Prophylactic and therapeutic HIV-1 vaccination strategies have been developed to prevent HIV-1 infection or for HIV-1 cure (1, 2). However, it is not clear how people with HIV-1 (PWH) under ART respond to different adjuvants and importantly, whether immune activation by adjuvants affects the so-called HIV-1 reservoir (3). Moreover, general vaccinations against influenza or other viruses are also cornerstones of preventative care in PWH (4–6).

Adjuvants are important components of vaccines to enhance immunogenicity. They activate innate pattern-recognition receptors on antigen presenting cells (APCs), provoking robust immune activation to improve the ensuing antigen-specific response (7, 8). Monophosphoryl lipid A (MPLA) is a toll-like receptor (TLR)4-targeting adjuvant; isolated from the lipopolysaccharide (LPS) of *Salmonella minnesota* R595, but modified to reduce toxicity and is sensed by immune cells through the TLR4 receptor (9). MPLA has been incorporated into approved vaccines both as a standalone adjuvant (MPL®) (10) and in combination with other components such as AS01b in the Shingrix (11) and AS04 in the Fendrix^TM12^ and Cervarix^TM^ vaccines (12) and TLR7 ligands (13). Subsequent studies have examined MPLA’s utility in HIV-1 vaccine candidates (14, 15), including its combination with saponin-based immune-stimulating complexes (ISCOMs). Saponins, derived from Quillaja saponaria, are used to form ISCOMs, which are cage-like 3D particles (∼40 nm in diameter) composed of saponin, cholesterol, and phospholipids (16–18). These ISCOMs promote B cell and T cell responses, in part by enhancing antigen delivery and uptake by APC (19, 20). One of these ISCOMs combines saponin (like QS-21) and MPLA into a Saponin-MPLA Nanoparticle (SMNP), by combining TLR4 stimulation (via MPLA) with inflammasome activation (via saponin), SMNP achieves more potent immune activation than MPLA in isolation (21).

While these adjuvants have been studied primarily for boosting adaptive immunity, their innate activation profiles in PWH on ART and their potential to influence the inducible HIV-1 reservoir remain insufficiently characterized. Here, we investigated whether TLR4-targeted adjuvants MPLA and SMNP induce strong cytokine responses in dendritic cells (DCs) and peripheral blood mononuclear cells (PBMCs) from healthy donors and from ART-suppressed PWH. Both SMNP and MPLA induced robust, dose-dependent activation of DCs. SMNP but not MPLA induced a strong cytokine response in PBMC from healthy donors, whereas this response was reduced in PBMC from ART-suppressed PWH. Notably, SMNP, but not MPLA, substantially reduced the size of the inducible HIV-1 reservoir in PWH *ex-vivo* without affecting bystander cells. Our findings indicate that while TLR4-targeted adjuvants can effectively induce immune activation, their potency appears reduced in ART-suppressed PWH, underscoring the need to develop adjuvants capable of overcoming HIV-associated immune dysfunction. As reservoir modulation was observed *ex vivo*, further evaluation is warranted to ensure that such adjuvants can be safely and effectively applied in therapeutic vaccination settings.

## Materials and Methods

### Study participants

The Amsterdam Cohort Studies (ACS) on HIV infection and AIDS is a prospective study among men who have sex with men and injecting drug users that started in 1984 (22). Nine HIV-1 infected participants who were on suppressive ART for at least 14 months were selected for this study (Table 1). All participants had an undetectable viral load for at least 11 months and their CD4 +T cell count recovered to ≥320 cells/mm3.

**Table 1.** Baseline characteristics of PWH.

| Characteristic | Value |
| --- | --- |
| Total participants | 9 |
| Sex: male | 9 |
| Age at time of sampling (median (years), range) | 40<sup>†</sup> (36-65) |
| CD4+ T cell count nadir (cells/mm<sup>3</sup>, range) | 327 (30–890) |
| Plasma viral load before ART initiation (copies/mL, range) | 184897 (13000–780000) |
| Time on effective ART (months, range) | 17.02 (14.04-25.28) |
| Duration of viral suppression (months, range)* | 14.32 (11.28-19.30) |
| CD4+ T cell count at sampling (cells/mm<sup>3</sup>, range) | 628 (320-1220) |
| Start ART (year, range) | 1997 (1996-2001) |
| ART regimen |  |
| Combination of (N)NRTI and PI | 1 |
| Combination of (N)NRTI | 8 |

<sup>†</sup>Months between the time point of first undetectable viral load and the sample analyzed.

### Ethics statement

Material from healthy controls was obtained from Sanquin, approval for which was given by the Ethics Advisory Body of the Sanquin Blood Supply Foundation in Amsterdam.

The ACS has been conducted in accordance with the ethical principles set out in the declaration of Helsinki. The study was approved by the institutional review board of the Academic Medical Center. Written informed consent was obtained from all participants (MEC 07–182; 20 August 2007). Data of the ACS were accessed for retrospective sample selection on December 18^th^, 2024 and sample selection was approved on December 20^th^ 2024. Authors had no access to information that could identify individual participants during or after data collection.

### Cell culture

PBMCs were obtained from buffy coats of healthy donors (Sanquin). PBMCs were isolated by a Lymphoprep (Axis-shield) gradient. PBMCs were cultured in Roswell Park Memorial Institute medium (RPMI, Thermo Fisher Scientific, Gibco) enriched with 10% FCS (Biological Industries), 10 IU/mL penicillin (Thermo Fisher), 10mg/mL streptomycin (Thermo Fisher), 2mM L-glutamine (Lonza) and 10IU/mL IL-2 (Invivogen) at 37°C in a humidified 5% CO2 incubator. Cells were stimulated with MPLA, SMNP or a LPS positive control for 24 hours after which supernatant was harvested for ELISA.

DCs were generated from PBMCs isolated from buffy coats of healthy donors. Monocytes were isolated by a Percoll (Amersham biosciences) gradient step. Immature monocyte-derived DCs were cultured for 6-7 days from monocytes in the presence of RPMI medium enriched with 10% FCS (Biological Industries), 10 IU/mL penicillin (Thermo Fisher), 10mg/mL streptomycin (Thermo Fisher), 2mM L-glutamine (Lonza), IL-4 (500U/mL, bioscource) and GM-CSF (800U/mL, invivogen) at 37°C in a humidified 5% CO2 incubator. Cells were stimulated with MPLA, SMNP or a LPS positive control for 24 hours after which supernatant was harvested for ELISA.

PBMCs from PWH obtained from the Amsterdam Cohort Studies were cultured in IMDM supplemented with 10% FCS, antibiotics (100 U/mL penicillin, 100 µg/mL streptomycin and ciproxine 5 µg/mL) and 20 U/mL IL-2 at 37°C in a humidified 5% CO2 incubator.

TZM-BL cells (RRID:CVCL_B478), which contain an HIV-1 LTR driven luciferase gene (23), were obtained through the NIH HIV Reagent Program, Division of AIDS, NIAID, NIH, and were cultured in Iscove’s Modified Dulbecco’s Medium (IMDM; Thermo Fisher Scientific, Gibco) supplemented with 10% fetal calf serum (FCS; HyClone, Cytiva, Marlborough, MA, USA) and antibiotics (100U/ml penicillin and 100ug/ml streptomycin) (Invitrogen, Carlsbad, CA, USA) at 37°C in a humidified 10% CO2 incubator.

J-Lat cells (clone A1) (RRID:CVCL_1G42) were obtained through the NIH HIV Reagent Program, Division of AIDS, NIAID, NIH. The J-Lat Tat-GFP clone A1 is a Jurkat cell harbouring an integrated HIV-1 LTR driving Tat and GFP expression. J-Lat A1 cells were cultured in IMDM supplemented with 10% FCS and antibiotics (100U/ml penicillin and 100ug/ml streptomycin) and exposed to DC supernatant from SMNP- or MPLA-treated conditions and fresh medium, in a 1:1 ratio. Two days later, cells were fixed and GFP expression and viability were analysed by flow cytometry to assess HIV-1 reactivation. Viability was determined by FCS/SSC. Conditions where cell death occurred were not included in the analysis. HEK293T (ATCC CRL-11268) (RRID:CVCL_0063) were cultured in IMDM (Thermo Fisher Scientific, Gibco) supplemented with 10% FCS L-glutamine and 1% penicillin/streptomycin. Cultures were maintained at 37°C in a humidified 5% CO2 incubator. HEK293T cells were transfected with TLR4 cDNA (HEK293T/TLR4) and were obtained through Dr. T Golenbock (24).

### Stimuli

MPLA liposomes were obtained through Darrell Irvine at a concentration of 1mg/mL and stored in suspension at 4°C.

SMNP adjuvant was obtained through Darrell Irvine at a concentration of 2.7mg/mL and stored in suspension at 4°C. The average particle size was 45nm.

LPS from Salmonella typhi (Sigma) was used at a concentration of 10 ng/mL.

### ELISA

Supernatant of MoDCs and PBMCs was harvested after 24 hours stimulation. Subsequently secretion of TNFα, IL-6, IL-10 and IL-12p70 proteins were measured by ELISA as described by the manufacturer (eBiosciences). OD450nm values were measured using BioTek synergy HT.

### Inducible HIV-1 reservoir reduction assay (HIVRRA)

The frequency of inducible HIV-1 infected CD4+ T cells from chronic PWH was determined by the HIVRRA assay (25, 26). In summary, PBMCs from PWH were exposed to 30µg/ml SMNP or 30µg MPLA directly or supernatant from DCs stimulated with 30µg/ml SMNP or 30µg MPLA for two days in the presence of 10μM Saquinavir. Following this, PBMCs were stimulated with 1μg/ml PHA in the presence of Saquinavir. After two days, PBMCs were washed and seeded in an 11-fold titration containing 3×10^4^ PBMCs in the first row and serially diluted 1:2 across a 96-well plate onto 2×10^4^ TZM-BL cells. After four days, the luciferase activity, driven by the HIV-1 LTR, was quantified in TZM-BL cells co-cultured with PBMCs from PWH by addition of 25 μl luciferase activity reagent (LAR) substrate (0.83mM ATP, 0.83 mM of d-Luciferin (Duchefa Biochemie B.V., Haarlem, The Netherlands), 18.7mM MgCl2, 0.78μM Na2H2P2O7, 38.9mM Tris (pH 7.8), 0.39% glycerol, 0.03% Triton X-100 and 2.6 μM dithiothreitol) and measuring luminescence in relative light units (RLU) using a luminometer (Berthold Technologies, Germany). The calculation of relative infectious units per million cells (IUPM) was based on 30% of the maximum RLU using logistic regression and is a measure for the functional HIV-1 reservoir.

To evaluate cytotoxicity of the compounds during the assay, a consistent amount of Cell Trace Violet (CTV)-labelled PBMCs from blood donors were used as spike-in and the ratio between CTV-labelled PBMC and unlabelled PWH PBMC was determined by flow cytometry. This also allowed to adjust for PWH PBMC cell input for subsequent reservoir calculations.

### Flow cytometry

Following 24h stimulation DCs were washed with PBS, then stained at 4C in the dark for 30 minutes with FITC conjugated anti-CD86 (1:25, Biolegend, San Diego, CA, USA, RRID AB_2721574), PE conjugated anti-CD80 (1:12.5, Biolegend, RRID AB_2890803), APC conjugated anti-CD83 (1:25 Biolegend, RRID AB_314519). After staining, cells were washed twice using FACS-PBA and analysed on a BD Symphony A1.

Healthy control PBMCs utilized in the HIVRRA were cultured as described in ‘cell culture’, then stained with CellTrace™ Violet (Thermo Fisher Scientific) according to the manufacturer’s protocol and fixed using FluoroFix™ Buffer (Biolegend).

J-lat cells were fixed using FluoroFix™ Buffer before GFP was measured.

## Statistical analysis

Statistical analysis of obtained data was performed using Graphpad Prism 10 (Graphpad Software Inc). Two-way ANOVA tests were performed to compare data between donors when triplicates were obtained. One-way ANOVA or Brown-Forsythe and Welch ANOVA tests were performed data between donors for data obtained in monoplo. Post-hoc corrections were performed. Differences were considered statistical significant when p < 0.05.

EC50 values were calculated by normalizing cytokine values obtained from donors to those of the LPS condition, setting the LPS condition to 100. Subsequently a logistic regression was performed at EC50 values were extracted.

## Results

### SMNP and MPLA activate DCs and induce pro-inflammatory cytokines

To evaluate the impact of SMNP and MPLA on DC activation, we first stimulated monocyte-derived DCs (DCs) from blood bank donors with increasing doses of each adjuvant and quantified the expression of T-cell priming proteins CD80, CD83, and CD86 by flow cytometry. Corresponding doses of MPLA and saponin QS-21 in each SMNP dose are shown in table 2.

**Table 2.** Contents of MPLA and QS-21 utilized in SMNP doses.

| SMNP | Content |  |
| --- | --- | --- |
| Dose<br>(µg/mL) | MPLA<br>(µg/mL) | QS-21<br>(µg/mL) |
| 0.01 | 0.00033 | 0.00246 |
| 0.05 | 0.00163 | 0.01230 |
| 0.1 | 0.00326 | 0.02459 |
| 0.5 | 0.01630 | 0.12 |
| 1 | 0.03259 | 0.25 |
| 5 | 0.16 | 1.23 |
| **10** | 0.33 | 2.46 |
| **20** | 0.65 | 4.92 |
| **30** | 0.98 | 7.38 |
| **50** | 1.63 | 12.30 |

Both SMNP and MPLA induced a dose-dependent upregulation of CD80, CD83 and CD86, comparable to LPS at high concentrations (Figure 1A-C). Notably, while both SMNP and MPLA reached a similar plateau of CD80, CD83 and CD86 expression at high concentration, SMNP stimulation led to upregulation of all markers at lower concentrations, with EC50 values for SMNP for CD80, CD83 and CD86 being 0.65, 1.82 and 0.65 μg/mL respectively (S1A-C Figure and S1 Table). Comparatively, for MPLA these were 2.27, 12.67 and 2.80 μg/mL (S1A-C Figure and S1 Table).

**Figure 1:**
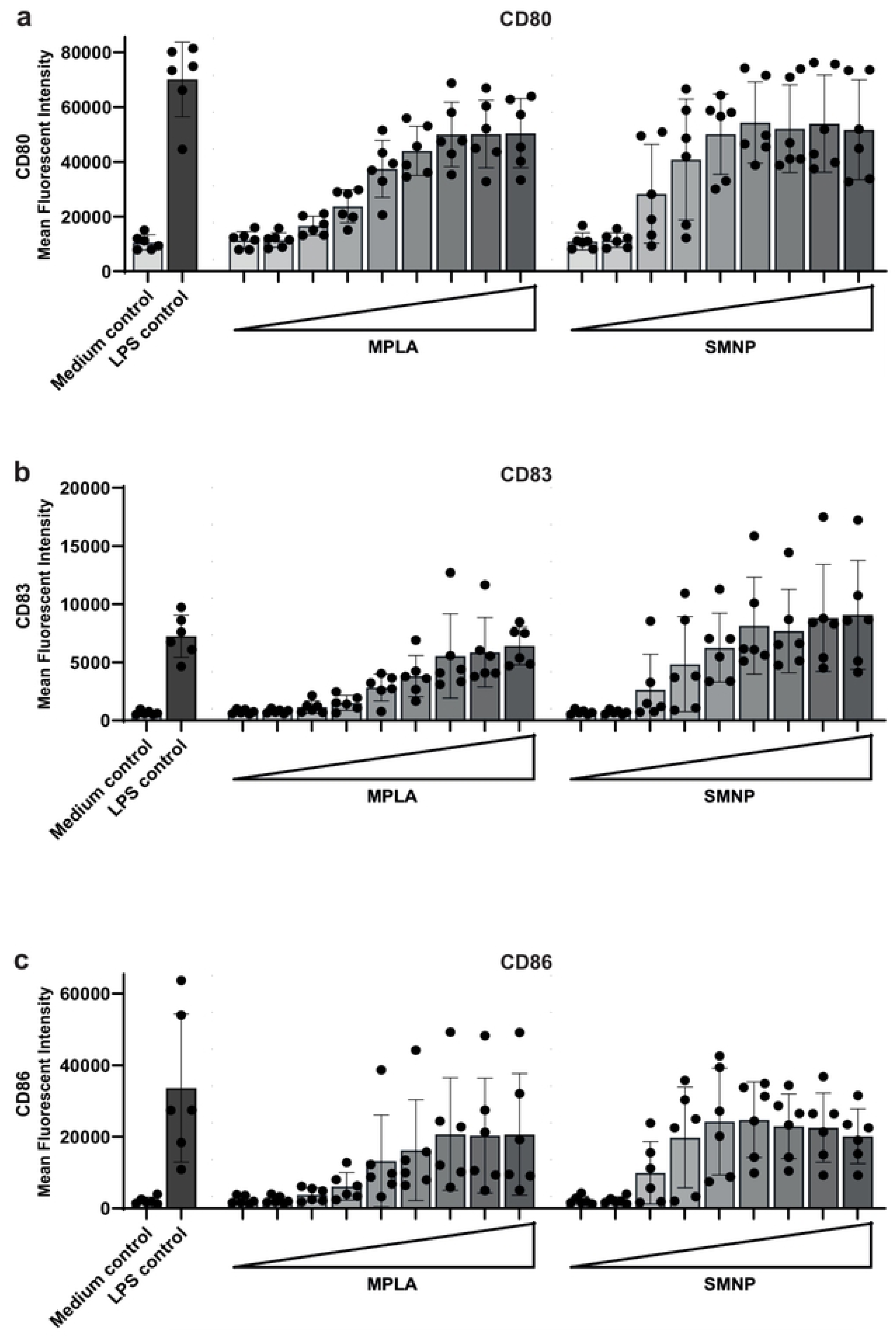
SMNP and MPLA activate DCs in a dose dependent manner (A-C) Surface expression of CD80, CD83 and CD86, was assessed on DCs from six different donors stimulated with increasing doses of SMNP and MPLA, (0.05 – 0.1 – 0.5 – 1 – 5 – 10 – 20 – 30 – 50μg/mL). A medium control and positive LPS (10ng/mL) control were added. DCs were stimulated for 24 hours and surface expression of co-stimulatory molecule CD80 (A), maturation marker CD83 (B) and co-stimulatory molecule CD86 (C) was measured using flow cytometry (n=6).

We next examined whether SMNP and MPLA differentially induce cytokine responses in DCs from blood bank donors. IL-6 and TNFα were selected as prototypical NF-κB–driven pro-inflammatory cytokines, IL-12p70 as a Th1-polarizing mediator, and IL-10 as a counter-regulatory cytokine that shapes the magnitude and quality of downstream responses. Both SMNP and MPLA triggered potent secretion of TNF-α, IL-6 and IL-12p70 in a dose dependent manner (Figure 2A, B, C). Similar to what we observed for co-stimulatory molecules, SMNP proved more potent than MPLA at inducting IL-6 and TNFα production, with SMNP’s EC50 being 1.00 μg/mL for IL-6 and 0.62 μg/mL for TNFα (S1D Figure, S1F Figure and S1 Table). Comparatively, for MPLA this was 2.15 μg/mL and 7.27 μg/mL, respectively (S1D Figure, S1F Figure and S1 Table).

**Figure 2:**
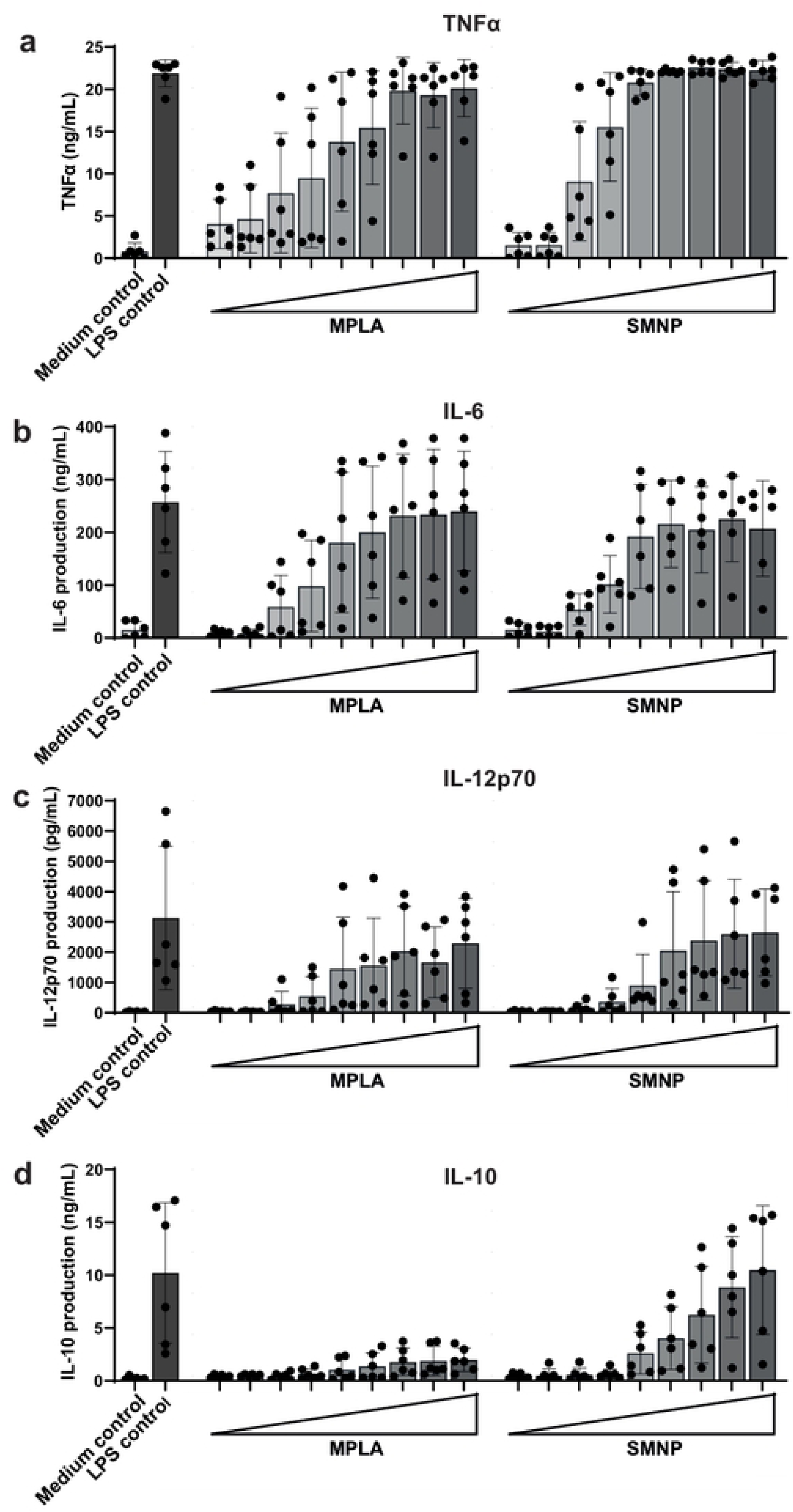
SMNP and MPLA induce pro-inflammatory cytokines in DCs. (A-D) DCs from six different donors were stimulated with SMNP and MPLA with increasing doses for 24 hours, (0.01 – 0.05 – 0.1 – 0.5 – 1 – 5 – 10 – 20 – 30 – 50μ g/mL) Subsequently, supernatant was harvested and the cytokines TNFα (A), IL-6 (B), IL-10 (C) and IL-12p70 (D) were measured using ELISA (n=6).

Notably, MPLA induced IL-12p70 at lower doses compared to SMNP, having an EC50 of 2.13 μg/mL compared to SMNP’s 9.83 μg/mL, with SMNP achieving a slightly higher plateau compared to MPLA at higher doses (Figure 2C, S1G Figure and S1 Table). Strikingly, IL-10 induction was much more potent following stimulation with SMNP compared to MPLA, with the higher concentrations reaching levels comparable to the LPS control (Figure 2D), reaching the EC50 at 20.65 μg/mL (S1E Figure and S1 Table).

### SMNP and MPLA trigger TLR4 to induce immune activation

MPLA, derived from LPS, is a known TLR4 agonist and SMNP contains MPLA but in a saponin cage that likely enhance binding to TLR4 (20, 27, 28). To assess whether SMNP induces TLR4 activation we utilized a TLR4-expressing Human Embryonic Kidney cells line (HEK293T/TLR4) (24). We quantified IL-8 as an NF-κB–dependent chemokine readout of TLR4 activation. After 24 hours stimulation of the parental HEK293T cells with MPLA, SMNP or the LPS control, no induction of IL-8 was observed. However, HEK293T/TLR4 cells responded to MPLA stimulation, albeit with lower levels of IL-8 being induced compared to SMNP. SMNP stimulation induced IL-8 secretion at comparable levels as LPS control (Figure 3). These data confirm that both MPLA and SMNP act as TLR4 agonists.

**Figure 3:**
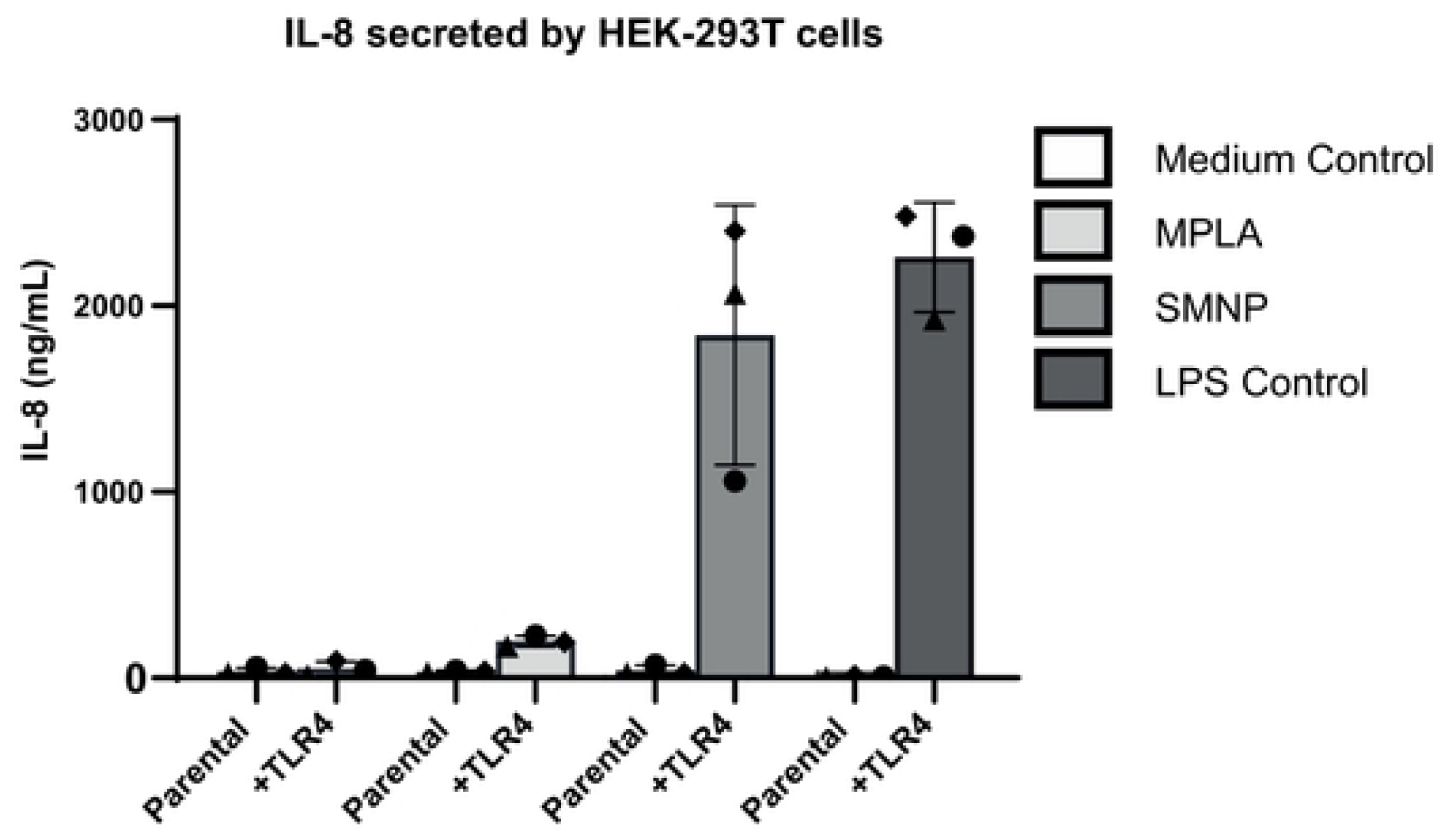
SMNP and MPLA induce TLR4 dependent immune activation HEK293T and HEK293T cells expressing recombinant TLR4 were stimulated with SMNP (30µg/mL), MPLA (30µg/mL) and an LPS (10ng/mL) positive control for 24h. Supernatant was harvested, and IL-8 secretion was measured with ELISA (n=3).

### Cytokines secreted by DCs reverse HIV-1 latency in J-Lat cells

As both MPLA and SMNP induced pro-inflammatory cytokines in DCs, we investigated whether supernatant from these stimulated DCs could reactivate HIV-1 using an HIV-1 latency model in J-Lat cells. In this model, J-lats harbour a Green Fluorescent Protein (GFP) under the control of the HIV-1 long terminal repeat (LTR). Following stimulation of DCs with MPLA and SMNP for 24 hours, the supernatant was harvested and transferred to the J-Lat cells. After 24 hours, latency reversal was determined by measuring GFP expression by flow cytometry. Supernatants from DCs stimulated with both MPLA and SMNP induced a strong dose-dependent increase in GFP+ J-Lat cells (Figure 4A). SMNP induced a higher overall re-activation compared to MPLA, closely approximating the response of the LPS positive control (Figure 4A). HIV-LTR activation occurred absent of cell-death (Figure 4B). These data suggest that both MPLA- and SMNP-induced immune activation leads to HIV-1 reactivation.

**Figure 4:**
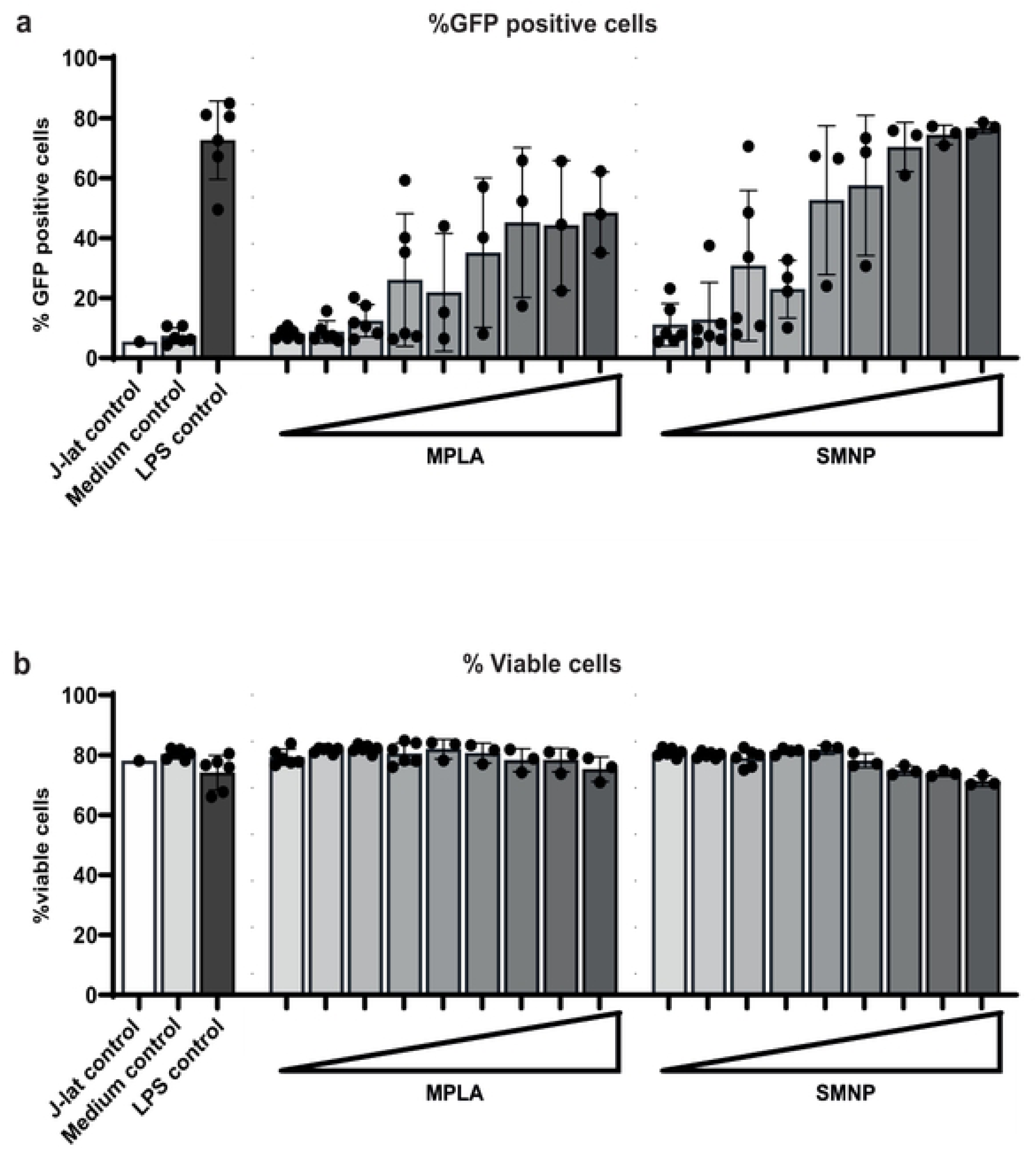
Adjuvants indirectly induce latency reversal in J-lat A1 cells. DCs were stimulated with SMNP and MPLA with increasing doses for 24 hours and supernatant was harvested. (A) J-lat cells were exposed to the supernatant and percentage of GFP positive cells was measured by flow cytometry after 24 hours of stimulation. (B) Only conditions with viable cells, as determined by FSC/SSC scattering relative to control conditions, were included in the analysis (n=6).

### SMNP, but not MPLA, induces cytokine production in PBMCs from PWH

Despite effective ART, the immune system of ART treated PWH have been shown to be dysfunctional compared to healthy donors (29–31). Here we investigated whether MPLA and SMNP induce immune activation in PWH (n=9), using PBMCs from PWH on ART (virologically supressed), and compared this to responses in PBMC from healthy donors. Baseline characteristics of PWH are displayed in table 2. SMNP and MPLA stimulation of PBMC from healthy donors led to detectable levels of IL-1β, TNFα, IL-6, IL-10 and IL-12, with SMNP being significantly more potent at inducing IL-1β, TNFα, IL-6, IL-10 than MPLA (Figure 5 and S2 Figure).

**Figure 5:**
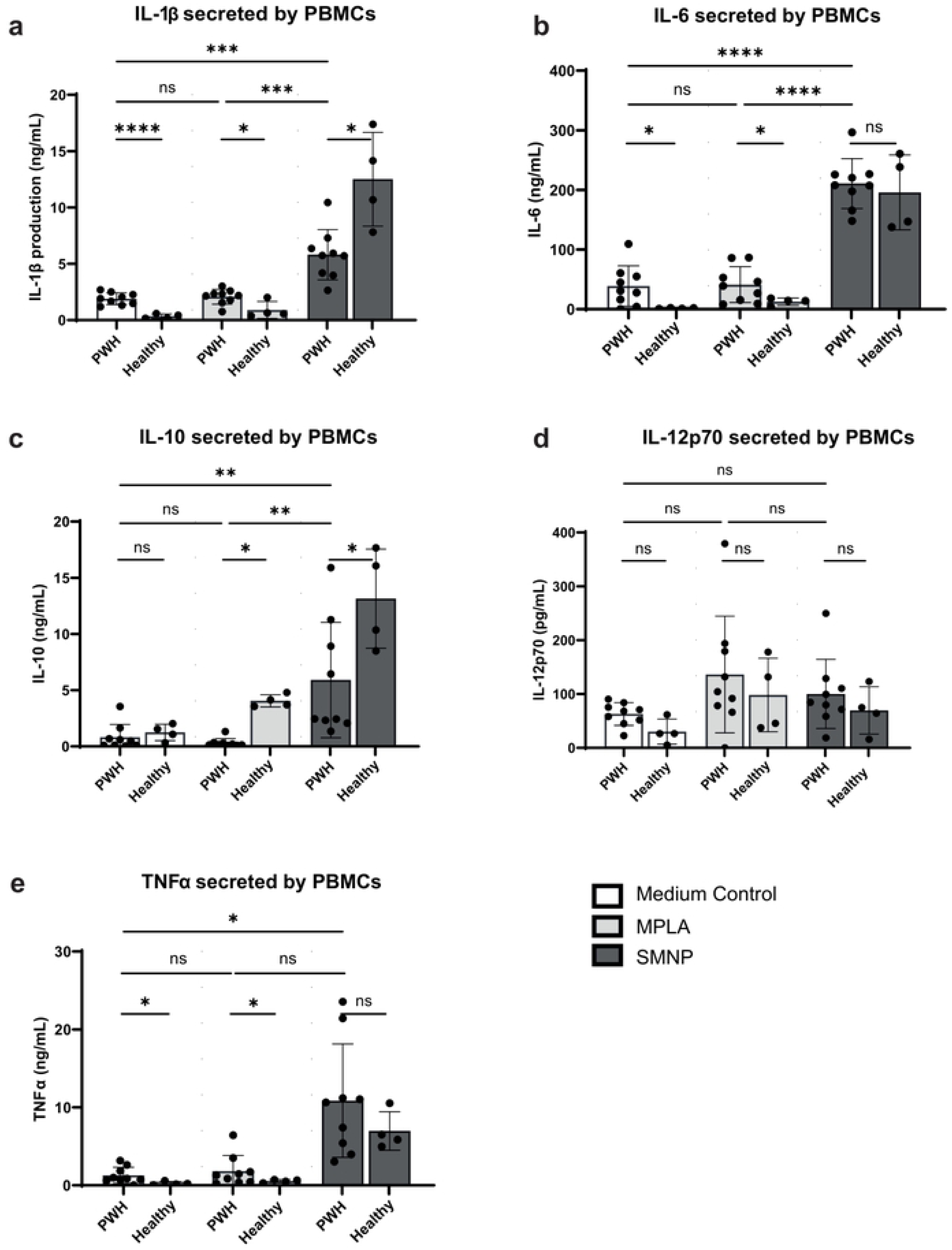
PBMCs from PWH induce similar levels of cytokines to SMNP and MPLA as in healthy donor controls. PBMCs from PWH (n=9) and healthy controls (n=4) were stimulated with SMNP (30µg/mL) and MPLA (30µg/mL) for 24 hours. Subsequently, supernatant was harvested and the cytokines IL-1β (A), IL-6 (B), IL-10 (C), IL-12p70 (D) and TNFα (E) were measured using ELISA. Comparisons between PWH and healthy donors were made using student Welch’s T-test (A-E). Comparisons within PWH were performed using a one-way ANOVA (A-C) or Brown-Forsythe and Welch ANOVA (D-E). Normality was tested for and controlled using QQ plots. Significant differences are indicated, *P < 0.05, **P < 0.01, ****P < 0.001, ****P < 0.0001.

In PWH, a similar trend was observed. Consistent with previous reports, the unstimulated controls showed a higher baseline level of cytokine induction, with significantly higher levels of IL-1β, IL-6, IL-12 and TNFα compared to healthy donors (Figure 5). Similarly, SMNP induced a potent induction of pro-inflammatory cytokines, encompassing IL-1β, TNFα, IL-6 and IL-10, while a nonsignificant increase in IL-12 was observed, which was also observed with MPLA. Interestingly, PBMCs from PWH showed differences in SMNP induced cytokine profile, with lower levels of IL-1β and IL-10 compared to healthy donors (Figure 5). MPLA did not lead to significant cytokine induction in PWH PBMCs, consistent with our observations in healthy donor PBMCs.

### SMNP, and to a lesser extent MPLA, reduces HIV-1 reservoirs in PWH *ex vivo*

We next investigated whether MPLA and SMNP can affect the HIV-1 reservoir in PBMCs from PWH *ex vivo*. To this end, PBMC from virologically supressed PWH (n=9) were stimulated with or without MPLA or SMNP, and after 24 hours the replication competent HIV-1 reservoir was measured using the inducible HIV-1 reservoir reduction assay (HIVRRA) (25, 26). In brief, PBMCs from PWH are activated using PHA and subsequently cocultured with TZM-BL reporter cells. Infectious virus produced by PBMCs from PWH is detected by luciferase activity (Figure 6C) and expressed as infectious units per million cells (IUPM).

**Figure 6:**
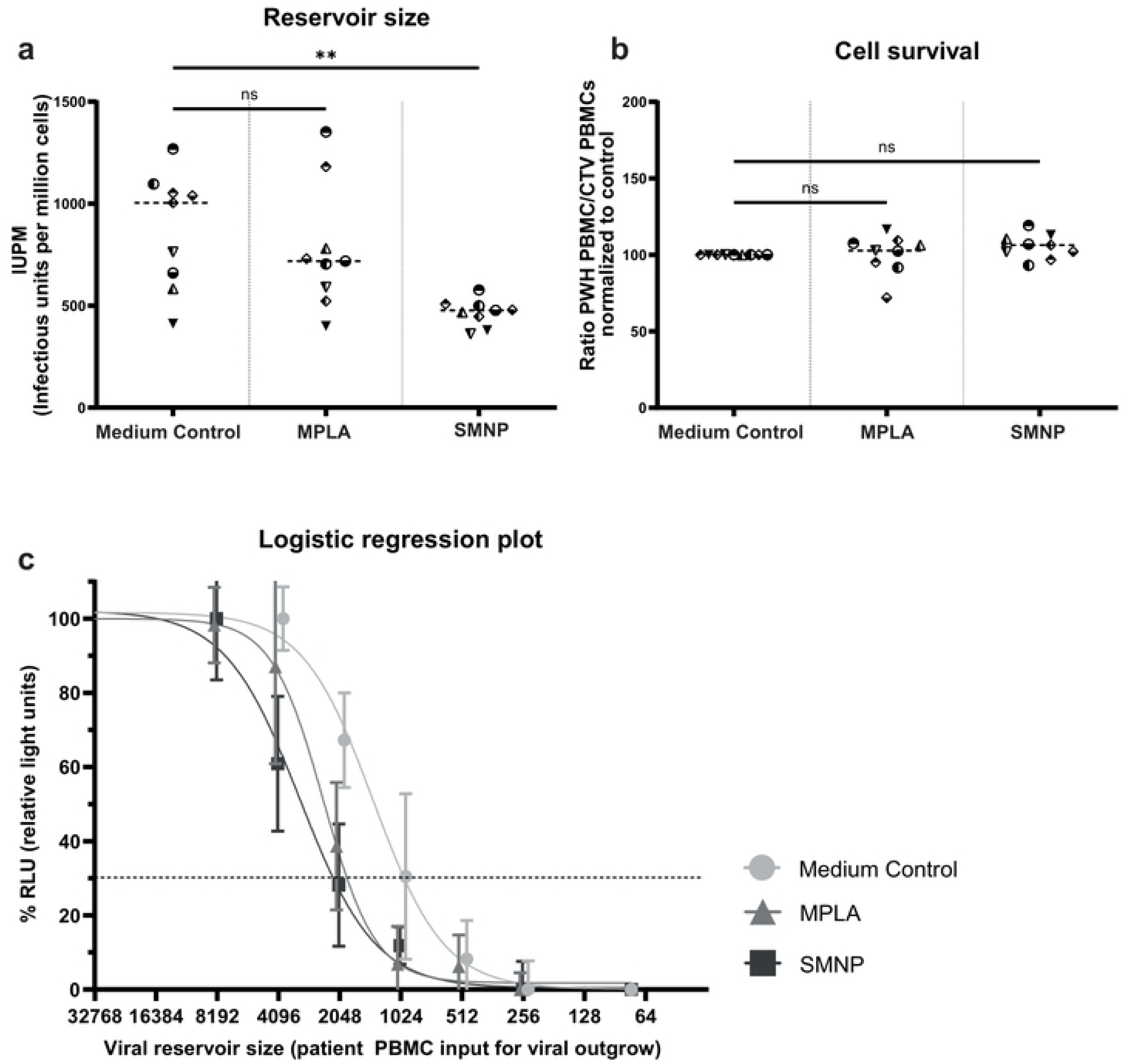
SMNP potently, and MPLA moderately, reduce HIV-1 reservoirs in PBMCs from PWH without inducing cytotoxicity as measured by the HIVRRA. (A) IUPM after treatment with the compounds for each participant (n=9); the median IUPM is displayed. Comparisons of each condition to control were made using a one way ANOVA. (B) Ratio of PWH PBMCs/healthy Cell trace violet (CTV)-stained PBMCs normalized to the medium control to check cytotoxicity of SMNP and MPLA. Comparisons of each condition to control were made using a Friedman test. (C) Logistic regression plot of the quantification of HIV-1-infected cells in PBMC from PWH after treatment with SMNP, MPLA or medium control using TZM-BL cells. Significant differences are indicated, **P < 0.01

SMNP exposure led to a significant reduction of the HIV-1 reservoir in PBMCs from PWH (Figure 6A). MPLA treatment did not lead to a significant reduction of the HIV-1 reservoir in PBMCs from PWH, although a reduction of the reservoir was observed in certain donors. Importantly, MPLA and SMNP treatment did not affect cell viability (Figure 6B), suggesting a preferential loss of infected cells upon MPLA and SMNP treatment rather than bystander toxicity.

To determine whether reduction of the inducible reservoir was a direct effect of the SMNP or MPLA stimulation or caused by the cytokine induction following SMNP or MPLA stimulation, PBMCs from PWH were stimulated with supernatant harvested from SMNP or MPLA stimulated DCs. Cytokine enriched supernatant from SMNP stimulated DCs showed a reduction in inducible reservoir compared to stimulation of PBMCs from PWH directly with SMNP (S3A Figure). PBMCs from PWH stimulated with MPLA or with supernatant from MPLA treated DCs, showed a minor decrease in inducible reservoir (S3A Figure). Notably, supernatant from adjuvant treated DCs did not affect bystander cells (S3B Figure).

## Discussion

Adjuvants are important to induce an efficient immune response against the antigens in vaccination. MPLA and its Saponin-caged derivative SMNP have been successfully used in pre-clinical vaccination studies (20, 27, 32). Consistent with their mechanism as TLR4 agonists, MPLA and SMNP triggered a broad immune response characterized by a surge in proinflammatory cytokines. We observed high levels of key cytokines produced by DCs shortly after stimulation, which is in line with the acute innate immune reaction typically seen with TLR4-agonists (28, 33). Indeed, we show that both adjuvants can induce a cytokine response through activation of TLR4. TLR4 engagement activates both MyD88- and TRIF-dependent signalling, leading to NF-κB and IRF activation and induction of inflammatory mediators (27). MPLA is a detoxified lipopolysaccharide derivative that acts as a TLR4 agonist with TRIF-biased signalling, preferentially triggering the TRIF-dependent pathway over the MyD88 pathway (27, 34). MPLA has potent immunostimulatory properties; it is a component of licensed adjuvant systems used in human vaccines (28, 35). However, responses to adjuvants require competent immunity. PWH have been shown to have chronic inflammation, a dysfunctional innate and adaptive immune response and a reduced response to vaccination(36–39). Here, we have investigated the efficacy of MPLA and SMNP to activate the immune response in healthy donors and PWH.

SMNP and MPLA induced a dose-dependent increase in activation markers and cytokine production in DCs from healthy donors, with SMNP but not MPLA also inducing IL-10. Notably, MPLA poorly induced cytokines in PBMCs compared to DCs, highlighting distinct responses between these cell populations. SMNP showed more consistent cytokine induction, inducing high levels of cytokines in both DCs and PBMCs. However, PWH showed a reduced cytokine response as compared to healthy donors. This dampened cytokine induction in PWH likely reflects underlying immune dysfunction associated with chronic HIV-1 infection even during effective ART (29, 40). Notably, immune exhaustion observed in PWH is associated with decreased potency of sentinel cells such as DCs or monocytes, as well as delayed and decreased responses to TLR agonists (41, 42). The lower IL-10 levels seen in PWH after SMNP stimulation might indicate an inability to mount the usual regulatory feedback or simply reflect the generally lower activation state. Since IL-10 is typically co-induced with IL-1β and TNFα during acute innate responses to limit pathology, its lack of proportional induction indicates possible dysregulation (43).

Pro-inflammatory cytokines, like TNFα, induce NF-κB signalling and drive HIV-1 gene expression (44). Our findings in the J-Lat latency model demonstrate that cytokines produced by MPLA and SMNP activated DCs can potently induce HIV-1 transcription. To extend these observations to PWH, we employed the HIVRRA to test the ability of MPLA and SMNP to reactivate, and subsequently reduce, the viral reservoir. We found that SMNP treatment led to a significant reduction in the inducible HIV-1 reservoir in PBMCs from virally suppressed PWH, whereas MPLA alone had no measurable impact. This lack of reservoir reduction with MPLA was not rescued by cytokine-rich supernatants from MPLA-stimulated DCs, suggesting that insufficient upstream signals beyond TLR4–TRIF bias may limit MPLA’s impact on the reservoir. SMNP mediated innate activation includes TLR4 priming and most likely QS-21 (saponin)–driven inflammasome activation, and this dual engagement of innate pathways is likely responsible for the observed differences between SMNP and MPLA.

TLR4 based activation, as induced by danger molecules like LPS, induce both early (MyD88) and late (TRIF) NF-κB activation, leading to the transcription of both NLRP3 (early) and pro-IL-1β (late), activating the inflammasome and leading to the induction of IL-1β through caspase 1 activity (27, 45). However, MPLA through removal of the phosphate group on Lipid-A, only activates late NF-κB (TRIF). The lack of a potent MyD88 signal modifies the fingerprint, typified by a reduced cytokine fingerprint (45). Together, TLR4 priming plus QS-21–driven inflammasome activation likely explains why SMNP elicits stronger cytokine responses and greater reservoir reduction than MPLA alone.

The use of TLR4 agonists within the HIVRRA assay illustrates that potent vaccine adjuvants can reactivate HIV-1 transcription through innate immune activation. In particular, SMNP stimulation of PBMCs from PWH resulted in a significant reduction of the inducible HIV-1 reservoir *ex vivo* without measurable bystander cytotoxicity, consistent with preferential effects on latently infected cells. TLR4 based activation relies on immune activation and transcription of pro-inflammatory cytokines (46).

Resting CD4+ T cells express minimal levels of TLR4, so the observed reservoir depletion likely stems from adjuvant-induced immune activation of antigen-presenting cells, leading to cytokine-mediated viral reactivation of latently infected T-cells. NF-κB in T-cells can be activated by cytokines, and promotes HIV-1 transcription since the LTR contains NF-κB binding sites (47, 48). The subsequent virus production may trigger cell death through cytopathic effects or apoptosis.

Interestingly, SMNP contains approximately 30-fold less MPLA than the MPLA-alone formulation, yet is markedly more potent, suggesting the saponin cage significantly enhances innate immune activation. Notably, saponin molecules (like QS-21, a component of SMNP) are known to directly activate the NLRP3 inflammasome in antigen-presenting cells. Saponins destabilize endolysosomal membranes, leading to potassium efflux and assembly of the NLRP3 inflammasome complex without involvement of MyD88 (48, 49). This event leads to caspase-1 activation, converting precursor cytokines like pro-IL-1β and pro-IL-18, induced by MPLA, into their biologically active forms. Strikingly, QS-21-driven inflammasome activation is markedly enhanced by prior TLR4-mediated transcriptional priming, which upregulates pro-IL-1β and inflammasome components (48, 50). Thus, MPLA provides the initial transcriptional priming, and QS-21 subsequently delivers the inflammasome-activating trigger, resulting in robust secretion of IL-1β (50).

Although most promising HIV vaccination strategies are deployed to induce broadly neutralizing antibodies to prevent HIV-1 acquisition (51–54), there is also great interests in therapeutic HIV-1 vaccine strategies in PWH aiming to reinforce virus-specific immunity to sustain virologic control off ART. For therapeutic vaccination strategies in PWH, HIV-associated immune dysregulation and altered TLR responsiveness should be taken into account (38, 55). In this context, our *ex vivo* data in ART-suppressed PWH show that SMNP has strong adjuvant activity, driving innate programs that support antigen presentation and T- and B-cell priming. However, SMNP induced responses also promote latency reversal with a reduction of the inducible reservoir without detectable bystander cytotoxicity in *ex vivo* assays. These properties position SMNP as a rational adjuvant for therapeutic HIV-1 vaccination in PWH. Importantly, clinical experience indicates that vaccination, including formulations with TLR4-based adjuvants, rarely cause clinically meaningful viremia in PWH on ART and does not compromise virologic control when ART is maintained (56–58).

Taken together, these observations support evaluating SMNP to strengthen vaccine responses in PWH and as a component of therapeutic vaccination strategies that pair immune potentiation with latency reversal; prospective clinical studies are warranted to determine whether these *ex vivo* effects translate into improved vaccine responses and measurable reservoir impact *in vivo*.

## Acknowledgements

We thank Darrell Irvine for kindly sharing SMNP.

The Amsterdam Cohort Studies on HIV infection and AIDS, a collaboration between the Public Health Service Amsterdam, the Amsterdam UMC of the University of Amsterdam, Medical Center Jan van Goyen and the HIV Focus Center of the DC-Clinics, are part of the Netherlands HIV Monitoring Foundation and financially supported by the Center for Infectious Disease Control of the Netherlands National Institute for Public Health and the Environment.

## Financial Disclosure

This research was funded by Health∼Holland/Aidsfonds(LSHM19101/P-44802) (TG/NK), Health∼Holland/Amsterdam UMC(2019-1167) (TG/NK), ZonMW/Aidsfonds(446002508) (TG/GdB), Aids-fonds/NWO (KICH2.V4P.AF23.001) (TG/NK/GdB/RS).

The funders had no role in study design, data collection and analysis, decision to publish, or preparation of the manuscript.

## Author contributions

Jade Jansen: Formal Analysis, Investigation, Methodology, Validation, Visualization, Writing – Original Draft Preparation, Writing – Review & Editing

Killian E. Vlaming: Formal Analysis, Investigation, Methodology, Validation, Visualization, Writing – Original Draft Preparation, Writing – Review & Editing

Leanne C. Helgers: Investigation, Methodology, Writing – Review & Editing Eva Schumacher: Resources, Writing – Review & Editing

Tanja M. Kaptein: Investigation, Writing – Review & Editing

Rogier W. Sanders: Resources, Supervision, Writing – Review & Editing

Godelieve J. de Bree: Resources, Conceptualization, Funding Acquisition, Supervision, Writing – Review & Editing

Teunis B.H. Geijtenbeek: Conceptualization, Funding Acquisition, Methodology, Project Administration, Supervision, Writing – Original Draft Preparation

Neeltje A. Kootstra: Conceptualization, Funding Acquisition, Methodology, Project Administration, Supervision, Writing – Original Draft Preparation

## Notes

### Competing Interest Statement

The authors have declared no competing interest.

